# Quantitative Measurements of Enlarged Perivascular Spaces in the Brain are Associated with Retinal Microvascular Parameters in Older Community-Dwelling Subjects

**DOI:** 10.1101/822155

**Authors:** Lucia Ballerini, Sarah McGrory, Maria del C. Valdés Hernández, Ruggiero Lovreglio, Enrico Pellegrini, Tom MacGillivray, Susana Muñoz Maniega, Ross Henderson, Adele Taylor, Mark E. Bastin, Fergus Doubal, Emanuele Trucco, Ian J. Deary, Joanna Wardlaw

## Abstract

**Background:** Perivascular Spaces (PVS) become increasingly visible with advancing age on brain MRI, yet their relationship to morphological changes in the underlying microvessels remains poorly understood. Retinal and cerebral microvessels share morphological and physiological properties. We compared computationally-derived PVS morphologies with retinal vessel morphologies in older people.

**Methods:** We analysed data from community-dwelling individuals who underwent multimodal brain MRI and retinal fundus camera imaging at mean age 72.55 years (SD=0.71). We assessed centrum semiovale PVS computationally to determine PVS total volume and count, and mean per-subject individual PVS length, width and size. We analysed retinal images using the VAMPIRE software suite, obtaining the Central Retinal Artery and Vein Equivalents (CRVE and CRAE), Arteriole-to-Venule ratio (AVR), and fractal dimension (FD) of both eyes. We investigated associations using general linear models, adjusted for age, gender, and major vascular risk factors.

**Results:** In 381 subjects with all measures, increasing total PVS volume and count were associated with decreased CRAE in the left eye (volume β=-0.170, count β=-0.184, p<0.001). No associations of PVS with CRVE were found. The PVS total volume, individual width and size increased with decreasing FD of the arterioles (a) and venules (v) of the left eye (total volume: FDa β=-0.137, FDv β=-0.139, p<0.01; width: FDa β=-0.144, FDv β=-0.158, p<0.01; size: FDa β=-0.157, FDv β=-0.162, p<0.01).

**Conclusions:** Increase in PVS number and size visible on MRI reflect arteriolar narrowing and lower retinal arteriole and venule branching complexity, both markers of impaired microvascular health. Computationally-derived PVS metrics may be an early indicator of failing vascular health and should be tested in longitudinal studies.

## 1. INTRODUCTION

Perivascular spaces (PVS) are also known also as Virchow-Robin spaces, and are seen with increasing clarity on brain MRI in older subjects, in small vessel disease (SVD), stroke, dementia and other neurological disorders [1]. PVS are fluid-filled compartments surrounding small perforating brain microvessels, mostly thought to be arterioles, and are thought to act as conduits for fluid transport, exchange between cerebrospinal fluid (CSF) and interstitial fluid (ISF) and clearance of waste products from the brain [2]. PVS are visible on T2w and T1w MRI when enlarged as thin linear or punctate structures (depending on scan orientation), oriented with perforating vessels, of similar signal to CSF [3, 4], having a diameter smaller than 3mm [5, 6]. PVS numbers have been reported to increase with age, with other brain SVD features [5], and with vascular risk factors, especially hypertension, in common brain disorders including stroke, mild cognitive impairment, and dementia including of vascular subtype (Debette et al., 2019, Francis et al., 2019).

To date, the quantification of enlarged PVS visible on MRI scans has mainly relied on qualitative ordinal visual scores [7]. Whilst shown to provide valuable information about PVS in aetiological studies to date, these scales are inherently insensitive to small details due to their limited number of categories, floor and ceiling effects, and may be affected by observer bias [7]. Computational tools for PVS quantification have been developed in the last five years [8-11]. The method by Ballerini *et al*. [11, 12] was able to segment PVS in the centrum semiovale and enabled quantification of several PVS features including the total count and total volume per individual subject’s brain, plus the size, length, width, shape and direction of each individual PVS. All these can then be analysed as mean or median per individual subject [11] or indeed per brain region. Previously, we showed good agreement between PVS visual rating and computational measures [11, 12]. We also showed associations between PVS individual widths and volume and WMH, which could indicate stagnation [13]. However it is not known if PVS morphologies reflect altered small vessel morphology.

The morphology of the retinal vessels is associated with stroke, including small vessel disease (SVD) (lacunar) stroke [14-17]. Arteriolar branching coefficients of retinal vessels were associated with white matter hyperintensities (WMH) in periventricular and deep white matter regions [18]. Fractal dimension, which reflects the complexity of the retinal vascular network, has been negatively related to WMH load and total SVD burden in older individuals [19]; and narrower retinal arterioles to enlarged perivascular spaces (PVS) seen on brain magnetic resonance imaging (MRI) [20]. Retinopathy has been associated with dementia, although associations with retinal vascular features are less clear[21].

Here we used retinal vessel measures as a surrogate for intracranial microvessel morphological alterations and tested for associations with computational PVS measurements to determine if changes in PVS morphology such as increased size or number were related to changes in small arterioles or venules indicative of vessel dysfunction.

## 2. MATERIALS AND METHODS

We used data from the Lothian Birth Cohort 1936 (LBC1936) Study [22]. The LBC1936 comprises 1091 community-dwelling individuals who took the Scottish Mental Health Survey in 1947, and gave informed consent to participate in this study at the age of 69. From the 866 study participants in the second wave of recruitment, at mean age 72.55 years (SD 0.71), 700 had structural MRI scans, and 814 had retinal scans of both eyes. Our analyses comprise data from those from which retinal and MRI data could be processed successfully.

The LBC1936 Study protocol was approved by the Lothian (REC 07/MRE00/58) and Scottish Multicentre (MREC/01/0/56) Research Ethics Committees (http://www.lothianbirthcohort.ed.ac.uk/).

All clinical and imaging acquisition methods and the visual and computational assessment of WMH and PVS visual scores in this cohort have been reported previously [22-24]. Briefly, medical history variables (hypertension, diabetes, hypercholesterolemia, cardiovascular disease history (CVD), smoking and stroke) were assessed at the same age as brain imaging. A history of CVD included ischaemic heart disease, heart failure, valvular heart disease and atrial fibrillation. Stroke data included clinically diagnosed stroke and also those with any ischaemic or haemorrhagic stroke seen in MRI scans in subjects with no clinical history of stroke. All medical history variables were coded as binary variables, indicating presence (1) or absence (0).

Structural brain MRI data were acquired using a 1.5-Tesla GE Signa Horizon HDx scanner (General Electric, Milwaukee, WI), with coronal T1-weighted (T1w), and axial T2-weighted (T2w), T2*-weighted (T2*w) and fluid-attenuated inversion recovery (FLAIR)-weighted whole-brain imaging sequences; details in [23].

Intracranial Volume (ICV) and WMH volume were measured on T2w, T1w, T1*w and FLAIR scans using a validated semi-automatic pipeline published in full previously [25]. For this study we express WMH as percentage of ICV.

The computational assessment of PVS used the T2w images acquired with 11,320ms repetition time, 104.9ms echo time, 20.83 KHz bandwidth, 2mm slice thickness, and 256 × 256mm field-of-view. The images were reconstructed to a 256 × 256 x 80 matrix, 1mm in-plane resolution. PVS were segmented in the centrum semiovale with the method described in [11]. This method was able to assess PVS in two pilot-size independent older age cohorts (age 64–72 years): individuals with a clinical diagnosis of dementia (*n* = 20), and patients who previously had minor stroke (*n* = 48) [11], and has been thoroughly validated on a large sample (n=533) of the LBC1936 [26]. The PVS computational method uses the three-dimensional Frangi filter [27], which parameters are optimized as described in [11, 12], to enhance and capture the 3D geometrical shape of PVS. It then calculates the PVS count and volume using connected component analysis [28]. The PVS count was defined as the number of connected-component objects in the segmented images. The PVS total volume was the total number of voxels classified as PVS. Individual PVS features (size, length, width) were also measured using connected component analysis. ‘PVS size’ was defined as the volume of each individual PVS to avoid confusion with ‘PVS volume’ which was the total volume of all the PVS in an individual subject. PVS length and width were defined, respectively, as the length of the longest and second-longest elongation axes of the ellipsoid approximating the PVS (see Figure 1b). Mean, median, standard deviation and percentiles of length, width and size were calculated for each subject. Before statistical analysis, the segmented PVS binary masks, superimposed on the T2W images, were visually checked by a trained operator, and accepted or rejected blind to all other data. To reduce operator input, images were checked but not edited. Other sequences (FLAIR and T1w) were checked in case of ambiguity and cases with other lesions detected as PVS were excluded. Images were deemed acceptable if the computational method was able to detect a reasonable amount of visible PVS, and did not detect too many artefacts as PVS. A small amount (< 20%) of false positives and negatives was considered acceptable. An example of PVS segmentation is shown in Figure 1a.

**Figure 1.**
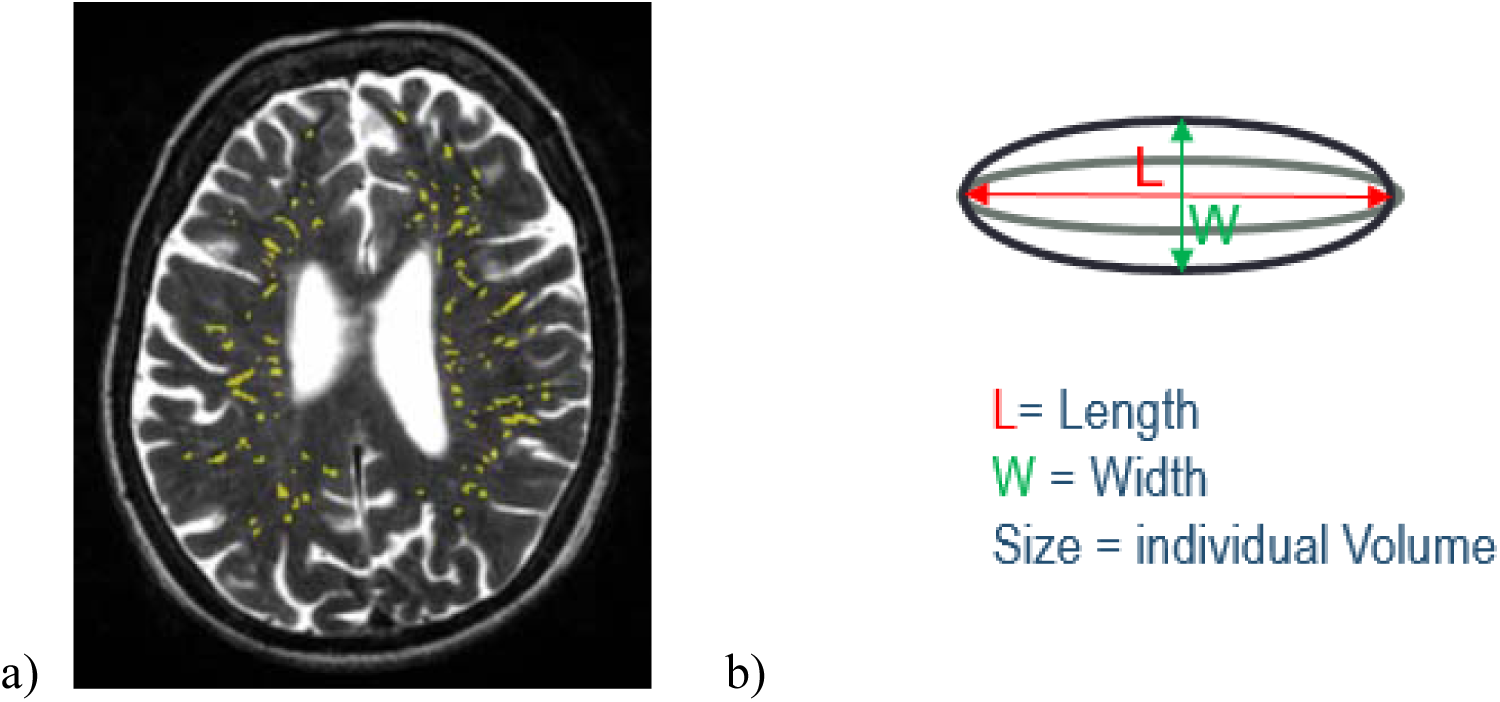
a) Axial T2-weighted slice showing in yellow the results of the PVS segmentation in a typical brain. b) Schematic illustration of the individual PVS metrics

Digital retinal fundus images were captured using a non-mydriatic camera at 45° field of view (CRDGi; Canon USA, Lake Success, New York, USA). All images were centred on the optic disc [29]. Retinal vascular measurements were computed for both eyes of each of the 601 included participants using the semi-automated software package, VAMPIRE (Vessel Assessment and Measurement Platform for Images of the REtina) version 3 [30, 31], by a trained operator blind to all other data (see the software interface in Figure 2). Quality assessment was performed by a trained operator. The reasons for image rejection included poor image quality, known pathologies or only one eye suitable for measurements. Briefly, the software detects the optic disc (OD) and the fovea in a retinal image. Next, the software detects the retinal blood vessels and classifies them as arterioles or venules. It then identifies the 6 major vessels of each type (artery, vein) in Zone B (a ring 0.5 to 1 OD diameters away from the centre) and measures their widths, combined in the summative coefficients described below. Details have been reported elsewhere [30, 31]. VAMPIRE 3.1 computes 151 morphometric measurements of the retinal vasculature, of which 5 were selected for this study: central retinal artery equivalent (CRAE), central retinal vein equivalent (CRVE), and arteriole-to-venule ratio (AVR), arteriolar and venular fractal dimension (FDa, FDv). CRAE, CRVE and AVR [32-34] were included as standard features summarizing the widths of major vessels near the optic disc. Based on recent findings on the associations between retinal features and brain imaging markers of SVD on this cohort [19], we also used FD.

**Figure 2.**
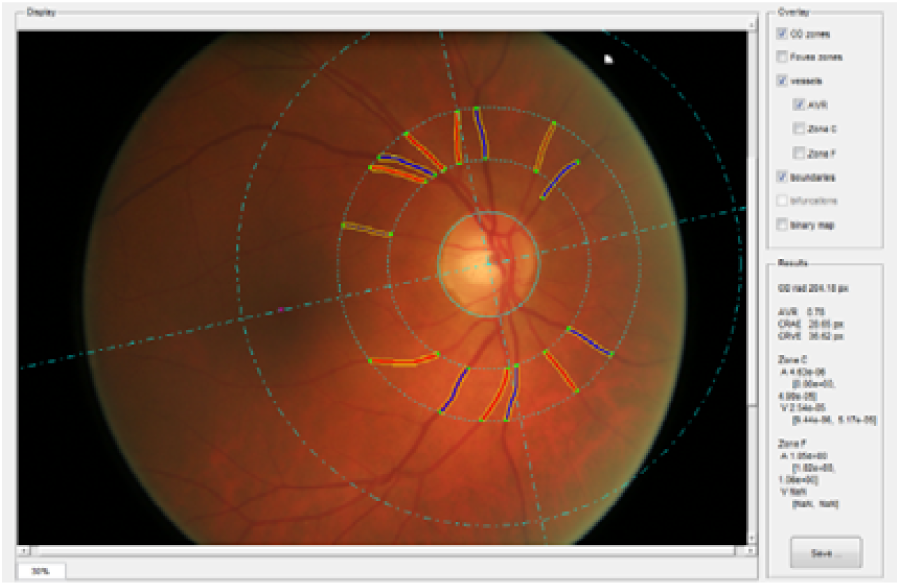
Retinal fundus image. Solid lines (red for arterioles and dark blue for venules) represent the vessels detected automatically and measured by VAMPIRE. Dotted lines represent the vessels widths

We included all subjects with usable PVS and retinal data from both eyes. Following recommendations from previous studies, and considering that high degrees of symmetry of any retinal or ocular structure, or the brain and major vessels to which it is connected, cannot be assumed [35], we analysed associations with each eye separately.

First, we compared the proportion of the sample for which all computational measures were available to those who were excluded using Welch two sample *t*-tests and chi-squared tests to identify any selection bias. Next, we generated descriptive statistics for all the variables involved in the analysis and calculated the bootstrapped bivariate cross-correlations between the retinal and brain variables, and controlled for false discovery rate (FDR). Finally, we estimated a series of generalized linear models to detect the associations between retinal vascular measures and PVS metrics. We adjusted for age, gender and vascular risk factors (hypertension, hypercholesterolemia, diabetes, cardiovascular disease, stroke and smoking).

## 3. RESULTS

PVS parameters were computed successfully in 540 out of 700 MRI images (77%). MRI scans that could not be processed successfully mainly contained motion artefacts that appeared as parallel lines similar to PVS. Fundus images of both eyes were available for 814 patients. Retinal measurements were computed successfully in both eyes for 601 out of 814 (74%) subjects. A total of 381 (44%) participants had both retinal and PVS measurements suitable for this study. Figure 3 shows a flowchart visualizing of the selection of our analytic sample.

**Figure 3.**
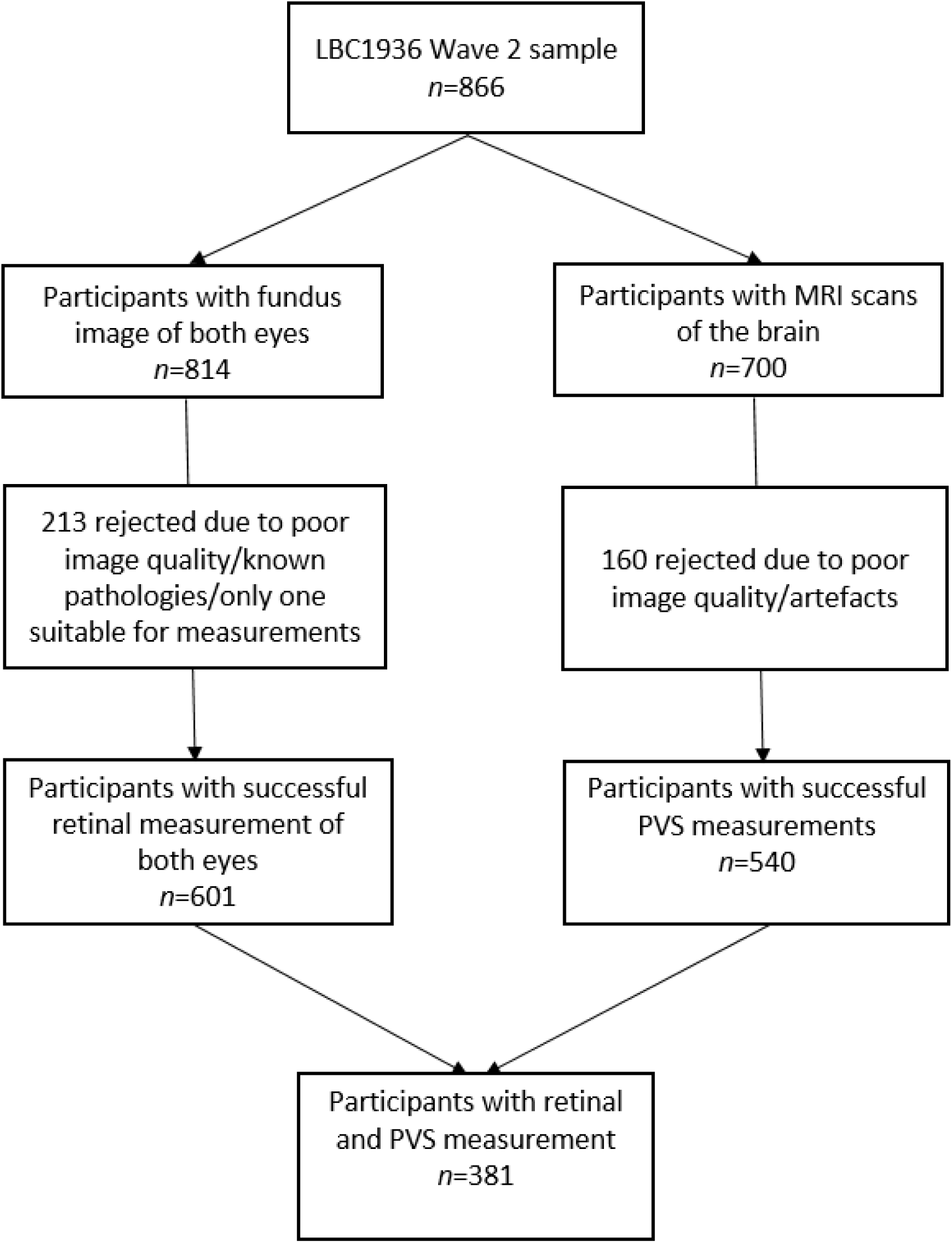
Flow chart of the analytic sample for the current study

Participants excluded from the retina-PVS analysis, due to missing either PVS or retinal data, did not differ from those included as to the proportion by gender, with diabetes, CVD and smokers (see Table 1). However, the group with successful computational PVS and retinal assessment were younger by an average of 44 days (included = 72.47 years, excluded = 72.60, p<0.01). The groups also differed in the proportion of patients with hypertension (included 43.31%, excluded 53.61%, p=0.003) and hypercholesterolaemia (included 34.91%, excluded 45.98%, p=0.001) indicating that the participants who were less likely to provide brain or retinal images that could be analysed computationally (due to movement eg) were older, with more co-morbidities.

**Table 1.**
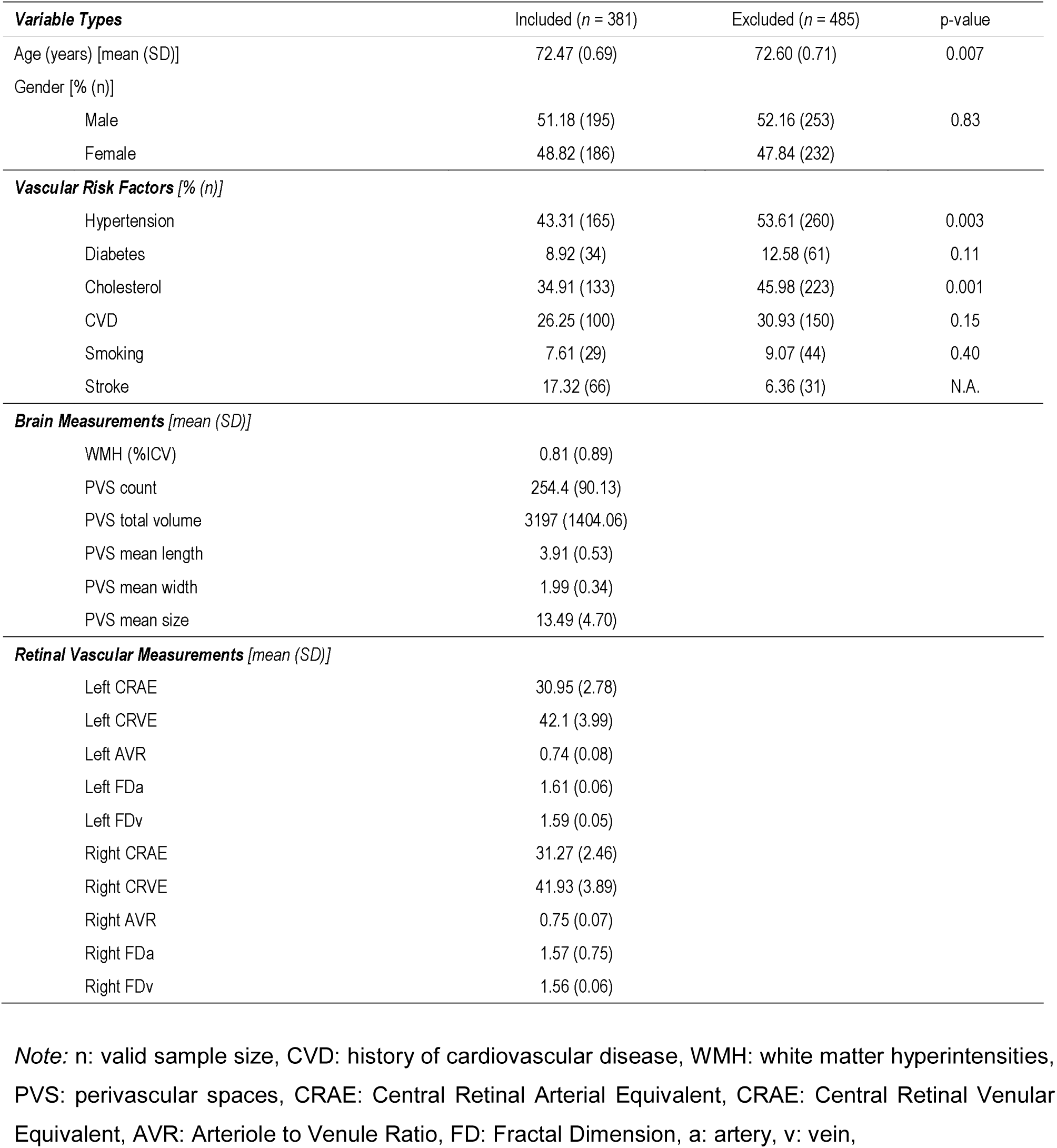
Brain imaging volumetric measures, vascular risk factors and retinal vascular measurements considered in the analyses and comparing those included vs excluded from the analysis.

| **Variable Types** | Included (*n* = 381) | Excluded (*n* = 485) | p-value |
| --- | --- | --- | --- |
| Age (years) [mean (SD)] | 72.47 (0.69) | 72.60 (0.71) | 0.007 |
| Gender [% (*n*)] |  |  |  |
| Male | 51.18 (195) | 52.16 (253) | 0.83 |
| Female | 48.82 (186) | 47.84 (232) |  |
| **Vascular Risk Factors** [% (*n*)] |  |  |  |
| Hypertension | 43.31 (165) | 53.61 (260) | 0.003 |
| Diabetes | 8.92 (34) | 12.58 (61) | 0.11 |
| Cholesterol | 34.91 (133) | 45.98 (223) | 0.001 |
| CVD | 26.25 (100) | 30.93 (150) | 0.15 |
| Smoking | 7.61 (29) | 9.07 (44) | 0.40 |
| Stroke | 17.32 (66) | 6.36 (31) | N.A. |
| **Brain Measurements** [mean (SD)] |  |  |  |
| WMH (%ICV) | 0.81 (0.89) |  |  |
| PVS count | 254.4 (90.13) |  |  |
| PVS total volume | 3197 (1404.06) |  |  |
| PVS mean length | 3.91 (0.53) |  |  |
| PVS mean width | 1.99 (0.34) |  |  |
| PVS mean size | 13.49 (4.70) |  |  |
| **Retinal Vascular Measurements** [mean (SD)] |  |  |  |
| Left CRAE | 30.95 (2.78) |  |  |
| Left CRVE | 42.1 (3.99) |  |  |
| Left AVR | 0.74 (0.08) |  |  |
| Left FD<sub>a</sub> | 1.61 (0.06) |  |  |
| Left FD<sub>v</sub> | 1.59 (0.05) |  |  |
| Right CRAE | 31.27 (2.46) |  |  |
| Right CRVE | 41.93 (3.89) |  |  |
| Right AVR | 0.75 (0.07) |  |  |
| Right FD<sub>a</sub> | 1.57 (0.75) |  |  |
| Right FD<sub>v</sub> | 1.56 (0.06) |  |  |

*Note:* *n*: valid sample size, CVD: history of cardiovascular disease, WMH: white matter hyperintensities, PVS: perivascular spaces, CRAE: Central Retinal Arterial Equivalent, CRAE: Central Retinal Venular Equivalent, AVR: Arteriole to Venule Ratio, FD: Fractal Dimension, a: artery, v: vein,

The 381 participants were of mean age 72.47 (SD 0.69), 186 (48.82%) were female, 165 (43.31%) had hypertension, 34 (8.92%) had diabetes, 133 (34.91%) were hypercholesteraemic, 100 (26.25%) had CVD, 29 (7.61%) were smokers, and 66 (17.32%) had prior clinical or imaging evidence of stroke (Table 1). The mean number of PVS in the centrum semiovale was 254.4 (SD 90.13). These correspond with median PVS visual ratings of 2, range 1-4 [7]. The mean percentage of WMH in the ICV was 0.81 (SD 0.89).

The cross-correlation matrix of bivariate associations among brain variables (PVS total volume, count, length, width, size) and retinal vascular measurements (CRAE, CRVE, AVR, FDa, FDv) is shown in Table 2. The PVS total volume and count were negatively correlated with left CRAE and AVR. The plots showing correlation among pairs of variable are shown in Figure 4, with histograms of the variables along the diagonal. No significant association was found between venular width and PVS. The total volume of PVS and the individual PVS size and width were negatively correlated with the arteriolar FD of the left eye. These results survived FDR correction. The direction of effect was similar for the right eye but did not reach significance (all p>0.01). The PVS measurements are all strongly associated with each other (Table 2). The correlations of retinal measures of the left-right eye are not high.

**Table 2.** Bivariate pairwise cross-correlations between the brain and retinal imaging variables evaluated. Spearman (ρ) values. Left eye values are given in the lower left part of the Table (light red) and Right values are given in the upper right part (light green). The significance level is indicated as follows: *p<0.05, **p<0.01 (Bold type). Results underlined survived FDR correction.

|  | PVS<br>total<br>volume | PVS<br>count | PVS<br>length | PVS<br>width | PVS<br>size | Right<br>CRAE | Right<br>CRVE | Right<br>AVR | Right<br>FDa | Right<br>FDv |
| --- | --- | --- | --- | --- | --- | --- | --- | --- | --- | --- |
| PVS<br>total<br>volume | 1 | **<u>0.924\*\*</u>** | **<u>0.868\*\*</u>** | **<u>0.875\*\*</u>** | **<u>0.852\*\*</u>** | <u>-0.123*</u> | -0.084 | -0.021 | -0.093 | -0.047 |
| PVS<br>count | **<u>0.924\*\*</u>** | 1 | **<u>0.710\*\*</u>** | **<u>0.686\*\*</u>** | **<u>0.627\*\*</u>** | -0.104* | -0.102* | 0.016 | -0.030 | -0.028 |
| PVS<br>length | **<u>0.868\*\*</u>** | **<u>0.710\*\*</u>** | 1 | **<u>0.939\*\*</u>** | **<u>0.922\*\*</u>** | <u>-0.129*</u> | -0.043 | -0.065 | <u>-0.118*</u> | -0.040 |
| PVS<br>width | **<u>0.875\*\*</u>** | **<u>0.686\*\*</u>** | **<u>0.939\*\*</u>** | 1 | **<u>0.971\*\*</u>** | <u>-0.128*</u> | -0.053 | -0.058 | <u>-0.129*</u> | -0.049 |
| PVS<br>size | **<u>0.852\*\*</u>** | **<u>0.627\*\*</u>** | **<u>0.922\*\*</u>** | **<u>0.971\*\*</u>** | 1 | <u>-0.127*</u> | -0.065 | -0.051 | <u>-0.145*</u> | -0.053 |
| Left CRAE | **<u>-0.177\*\*</u>** | **<u>-0.191\*\*</u>** | <u>-0.123*</u> | <u>-0.114*</u> | <u>-0.121*</u> | **<u>0.406\*\*</u>** | **<u>0.154\*\*</u>** | **<u>0.153\*\*</u>** | <u>0.123*</u> | 0.018 |
| Left CRVE | -0.035 | -0.047 | -0.026 | -0.038 | -0.037 | **<u>0.236\*\*</u>** | **<u>0.331\*\*</u>** | <u>-0.126*</u> | 0.110* | 0.112* |
| Left AVR | <u>-0.126*</u> | **<u>-0.133\*\*</u>** | -0.077 | -0.059 | -0.068 | **<u>0.139\*\*</u>** | **<u>-0.159\*\*</u>** | **<u>0.256\*\*</u>** | 0.015 | -0.093 |
| Left FDa | **<u>-0.134\*\*</u>** | -0.088 | <u>-0.120*</u> | **<u>-0.136\*\*</u>** | **<u>-0.150\*\*</u>** | **<u>0.176\*\*</u>** | **<u>0.242\*\*</u>** | -0.087 | **<u>0.307\*\*</u>** | **<u>0.285\*\*</u>** |
| Left FDv | <u>-0.119*</u> | -0.093 | -0.109* | <u>-0.127*</u> | **<u>-0.140\*\*</u>** | 0.034 | **<u>0.163\*\*</u>** | <u>-0.112*</u> | <u>0.127*</u> | **<u>0.273\*\*</u>** |

*Note:* PVS: perivascular spaces, CRAE: Central Retinal Arterial Equivalent, CRAE: Central Retinal Venular Equivalent, AVR: Arteriole to Venule Ratio, FD: Fractal Dimension, a: artery, v: vein.

**Figure 4.**
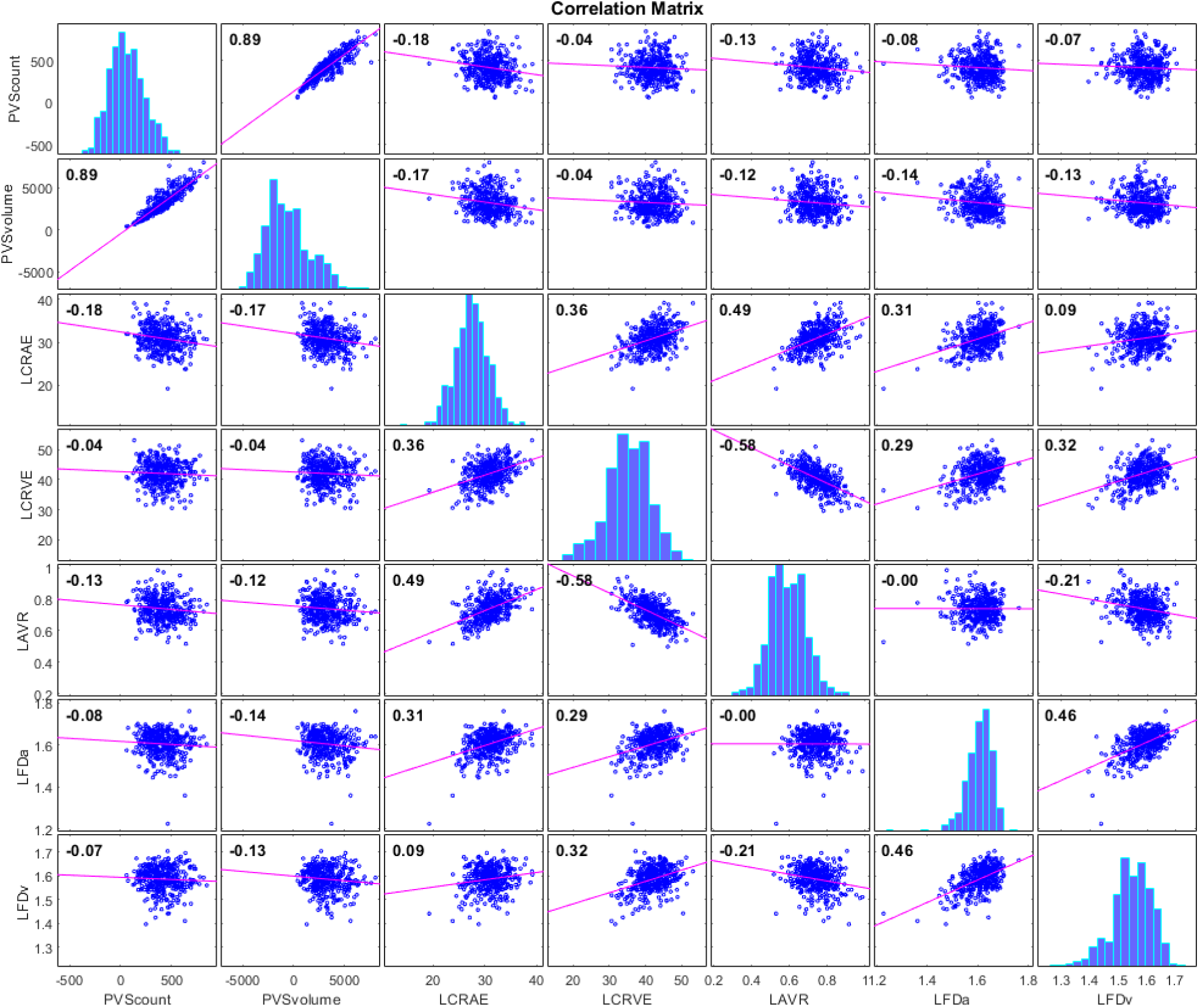
Correlation plots of count and total volume of PVS in the centrum semiovale vs left CRAE, CRVE, AVR, FDa and FDv. Histograms of the variables appear along the matrix diagonal.

We also tested for significant associations using general linear models (Table 3). After full covariate adjustment, the total volume and count of PVS were independently and negatively associated with left CRAE (volume β=-0.170, p=0.001; count β=-0.184, p<0.001). The association between the number of PVS with AVR attenuated to non-significance after adjustments. After correcting for covariates, the PVS total volume was associated negatively with the FD of the arterial and venular vasculature of the left eye (FDa β=-0.137, FDv β=-0.139, both p<0.01). Individual PVS width and size were both associated negatively with FD of the retinal arterioles and venules (width: FDa β=-0.144, FDv β=-0.158, both p<0.01; size: FDa β=-0.157, p=0.002; FDv β=-0.162, p=0.001). Smoking was also associated with increased individual PVS width and size (β range = 139 to160, all p<0.005). These associations survived FDR correction.

**Table 3.** Associations of brain and retinal imaging variables adjusted for age, gender and vascular risk factors. Bold type indicates p<0.05. Results underlined survived FDR correction.

| Model no. | Outcome name | Predictors (in addition to age and biological sex) |  |  |  |  |  |
| --- | --- | --- | --- | --- | --- | --- | --- |
|  |  | Name | Unstandardised values |  | Standardised values |  | p-value |
|  |  |  | B | SE | Beta | SE |  |
| 1 | **PVS total volume** | Hypertension | 1.59e-04 | 1.30e-04 | 0.066 | 0.055 | 0.22 |
|  |  | Diabetes | 1.98e-04 | 2.17e-04 | 0.052 | 0.057 | 0.36 |
|  |  | Hypercholesterolaemia | -1.58e-04 | 1.34e-04 | -0.065 | 0.055 | 0.24 |
|  |  | History of CVD | 4.53e-05 | 1.41e-04 | 0.017 | 0.053 | 0.75 |
|  |  | **Smoking** | **5.73e-04** | **2.28e-04** | **0.127** | **0.051** | **0.012** |
|  |  | Previous stroke | 2.95e-04 | 1.64e-04 | 0.093 | 0.052 | 0.07 |
|  |  | **Left CRAE** | **<u>-7.33e-05</u>** | **<u>2.21e-05</u>** | **<u>-0.170</u>** | **<u>0.051</u>** | **<u>0.00101</u>** |
| 2 | **PVS total volume** | Hypertension | 1.43e-04 | 1.31e-04 | 0.060 | 0.054 | 0.27 |
|  |  | Diabetes | 2.18e-04 | 2.18e-04 | 0.057 | 0.057 | 0.32 |
|  |  | Hypercholesterolaemia | -1.57e-04 | 1.35e-04 | -0.065 | 0.055 | 0.24 |
|  |  | History of CVD | 3.25e-05 | 1.41e-04 | 0.001 | 0.053 | 0.98 |
|  |  | **Smoking** | **5.27e-04** | **2.29e-04** | **0.117** | **0.051** | **0.022** |
|  |  | **Previous stroke** | **3.48e-04** | **1.65e-04** | **0.110** | **0.052** | **0.035** |
|  |  | **Left FDa** | **-2.89e-03** | **1.07e-03** | **-0.137** | **0.051** | **0.00704** |
| 3 | **PVS total volume** | Hypertension | 1.77e-04 | 1.31e-04 | 0.074 | 0.055 | 0.18 |
|  |  | Diabetes | 2.12e-04 | 2.18e-04 | 0.055 | 0.057 | 0.33 |
|  |  | Hypercholesterolaemia | -1.96e-04 | 1.36e-04 | -0.081 | 0.057 | 0.15 |
|  |  | History of CVD | -2.35e-07 | 1.41e-04 | -0.009 | 0.053 | 0.86 |
|  |  | **Smoking** | **5.53e-04** | **2.29e-04** | **0.123** | **0.051** | **0.016** |
|  |  | **Previous stroke** | **3.62e-04** | **1.65e-04** | **0.115** | **0.052** | **0.029** |
|  |  | **Left FDv** | **-3.32e-03** | **1.22e-03** | **-0.139** | **0.051** | **0.00678** |
| 4 | **PVS count** | Hypertension | 8.11 | 9.86 | 0.045 | 0.055 | 0.41 |
|  |  | Diabetes | 16.08 | 16.48 | 0.056 | 0.057 | 0.33 |
|  |  | Hypercholesterolaemia | -5.56 | 10.16 | -0.030 | 0.055 | 0.59 |
|  |  | History of CVD | 0.43 | 10.71 | 0.002 | 0.054 | 0.97 |
|  |  | Smoking | 18.41 | 17.29 | 0.054 | 0.051 | 0.29 |
|  |  | Previous stroke | 3.59 | 12.43 | 0.015 | 0.052 | 0.77 |
|  |  | **Left CRAE** | **<u>-5.95</u>** | **<u>1.68</u>** | **<u>-0.184</u>** | **<u>0.052</u>** | **<u>0.00043</u>** |
| 5 | PVS count | Hypertension | 5.44e+00 | 9.98e+00 | 0.030 | 0.055 | 0.59 |
|  |  | Diabetes | 1.71e+01 | 1.67e+01 | 0.059 | 0.058 | 0.31 |
|  |  | Hypercholesterolaemia | -4.16e+00 | 1.03e+01 | -0.023 | 0.056 | 0.69 |
|  |  | History of CVD | -7.74e-01 | 1.09e+01 | -0.004 | 0.051 | 0.94 |
|  |  | Smoking | 1.74e+01 | 1.75e+01 | 0.051 | 0.052 | 0.32 |
|  |  | Previous stroke | 5.61e+00 | 1.26e+01 | 0.024 | 0.053 | 0.66 |
|  |  | Left AVR | -1.26e+02 | 6.20e+01 | -0.105 | 0.052 | 0.0433 |
| 6 | **PVS width** | Hypertension | 5.54e-02 | 3.71e-02 | 0.081 | 0.054 | 0.14 |
|  |  | Diabetes | 3.69e-02 | 6.20e-02 | 0.034 | 0.057 | 0.55 |
|  |  | Hypercholesterolaemia | -5.99e-02 | 3.82e-02 | -0.086 | 0.055 | 0.11 |
|  |  | History of CVD | 6.84e-03 | 4.01e-02 | 0.009 | 0.053 | 0.86 |
|  |  | **Smoking** | **<u>1.80e-01</u>** | **<u>6.51e-02</u>** | **<u>0.139</u>** | **<u>0.051</u>** | **<u>0.00612</u>** |
|  |  | **Previous stroke** | **<u>1.10e-01</u>** | **<u>4.67e-02</u>** | **<u>0.122</u>** | **<u>0.052</u>** | **<u>0.019</u>** |
|  |  | **Left FDa** | **<u>-8.66e-01</u>** | **<u>3.03e-01</u>** | **<u>-0.144</u>** | **<u>0.050</u>** | **<u>0.00444</u>** |
| 7 | **PVS width** | Hypertension | 6.63e-02 | 3.71e-02 | 0.097 | 0.054 | 0.08 |
|  |  | Diabetes | 3.53e-02 | 6.19e-02 | 0.032 | 0.057 | 0.57 |
|  |  | Hypercholesterolaemia | -7.26e-02 | 3.84e-02 | -0.104 | 0.055 | 0.06 |
|  |  | History of CVD | -1.65e-03 | 4.00e-02 | -0.002 | 0.053 | 0.97 |
|  |  | **Smoking** | **<u>1.87e-01</u>** | **<u>6.49e-02</u>** | **<u>0.145</u>** | **<u>0.050</u>** | **<u>0.00415</u>** |
|  |  | **Previous stroke** | **<u>1.15e-01</u>** | **<u>4.67e-02</u>** | **<u>0.128</u>** | **<u>0.052</u>** | **<u>0.014</u>** |
|  |  | **Left FDv** | **<u>-1.08e+00</u>** | **<u>3.46e-01</u>** | **<u>-0.158</u>** | **<u>0.051</u>** | **<u>0.00197</u>** |
| 8 | **PVS size** | Hypertension | 5.73e-01 | 5.08e-01 | 0.061 | 0.054 | 0.26 |
|  |  | Diabetes | 1.18e-01 | 8.50e-01 | 0.008 | 0.057 | 0.89 |
|  |  | Hypercholesterolaemia | -6.18e-01 | 5.24e-01 | -0.065 | 0.055 | 0.24 |
|  |  | History of CVD | 7.81e-01 | 5.49e-01 | 0.008 | 0.053 | 0.89 |
|  |  | **Smoking** | **<u>2.73e+00</u>** | **<u>8.93e-01</u>** | **<u>0.154</u>** | **<u>0.050</u>** | **<u>0.00243</u>** |
|  |  | **Previous stroke** | **<u>1.50e+00</u>** | **<u>6.41e-01</u>** | **<u>0.121</u>** | **<u>0.052</u>** | **<u>0.019</u>** |
|  |  | **Left FDa** | **<u>-1.30e+01</u>** | **<u>4.15e+00</u>** | **<u>-0.157</u>** | **<u>0.050</u>** | **<u>0.00194</u>** |
| 9 | **PVS size** | Hypertension | 7.27e-01 | 5.10e-01 | 0.077 | 0.054 | 0.16 |
|  |  | Diabetes | 9.14e-02 | 8.50e-01 | 0.006 | 0.056 | 0.91 |
|  |  | Hypercholesterolaemia | -7.96e-01 | 5.27e-01 | -0.083 | 0.055 | 0.13 |
|  |  | History of CVD | -4.35e-02 | 5.50e-01 | -0.004 | 0.053 | 0.94 |
|  |  | **Smoking** | **<u>2.84e+00</u>** | **<u>8.91e-01</u>** | **<u>0.160</u>** | **<u>0.050</u>** | **<u>0.00155</u>** |
|  |  | **Previous stroke** | **<u>1.57e+00</u>** | **<u>6.42e-01</u>** | **<u>0.127</u>** | **<u>0.052</u>** | **<u>0.015</u>** |
|  |  | **Left FDv** | **<u>-1.52e+01</u>** | **<u>4.75e+00</u>** | **<u>-0.162</u>** | **<u>0.051</u>** | **<u>0.00149</u>** |

*Note:* PVS: perivascular spaces, CRAE: Central Retinal Arterial Equivalent, CRAE: Central Retinal Venular Equivalent, AVR: Arteriole to Venule Ratio, FD: Fractal Dimension, a: artery, v: vein, CVD: history of cardiovascular disease.

## 4. DISCUSSION

We found that increases in PVS total volume and count are associated with narrower retinal arterioles, and increased total volume, individual width and size of PVS are associated with decreased arteriolar and venular fractal dimension, independent of co-variates. Since both arteriole narrowing and reduced branching complexity are known indicators of adverse microvascular health, this work provides further evidence that an increased count of visible PVS and enlargement of individual PVS reflect underlying microvascular pathology rather than being an epiphenomenon.

To our best knowledge, this is the first study to compare multiple computational measures of PVS enlargement with retinal vascular measurements. Only one previous study reported associations between retinal vessel width and PVS using visual rating scales [20] and found that narrower arteriolar calibre, and to a lesser extent wider venular calibre, were significantly associated with higher numbers of PVS. The authors hypothesized that a failure in the CSF transmission may result in hemodynamic pressure differences that might manifest in changes in vascular calibre. They also hypothesized that narrower arterioles may lead to hypoperfusion, resulting in atrophy, and thus to PVS enlargement. Our negative associations between computational PVS metrics and arteriolar calibres are in the same direction. However, we did not find significant associations with venular calibres. Our findings support the hypothesis that increasing PVS total volume, count and individual size are markers primarily of arteriolar pathology in the brain. There is evidence of venular disease in ageing, SVD and dementia [36], but our results do not support a strong association of PVS and venular pathology.

A previous study, using this same cohort [19], reported associations between fractal dimension and WMH, but not PVS. However, this analysis used a visual rating scale and considered PVS in the basal ganglia, while our computational measurements, potentially more sensitive, are in the centrum semiovale. This might reflect that the vasculature differs between brain regions, possibly due to regional variations in the vessel and PVS anatomy, underpinning differences in distributions of fibrohyaline thickening, lipohyalinosis and cerebral amyloid angiopathy, which, in turn, may affect vessel-brain fluid exchange and PVS morphology [37-39].

The present findings support the hypothesis that retinal FD is a possible indicator of the state of health of the brain vasculature and indicate changes taking place in cerebral small vessels [19]. In line with our results, a previous study [40] reported that decreased retinal arterial FD was associated with cerebral microbleeds, while another study found that decreased FD was associated with small vessel (lacunar) versus non-small vessel ischaemic stroke [14]. It should however be kept in mind that the stability of the FD of the retinal and brain vasculature is under scrutiny [41, 42].

Several clinical and population-based studies have shown associations between retinal vascular changes and markers of cerebral SVD, as summarised by Hilal *et al*. [40], suggesting that changes in retinal vascular measures may be an early manifestation of cerebral SVD [43]. A systematic review and meta-analysis of associations between retinal vascular morphologies and dementia [21] found conflicting results for vessel calibre measurements, with the most consistent finding being a decreased fractal dimension in Alzheimer’s disease.

We found a significant association between smoking and PVS size and width in the models including fractal dimension. This is consistent with previous studies indicating the deleterious effects of smoking on brain structure [44, 45] and increased total burden of SVD [46]. However this conflicts with findings from the Three-City Dijon MRI study that did not find associations between smoking and PVS visual scores [47].

Eye laterality also deserves attention. Some studies choose to analyse the eye with the best image quality [20]; others use either the left or right eye [48]. We decided to analyse the associations with both eyes separately and found asymmetric results. This supports some of the conclusions of a laterality study [35]. The reasons why the morphology of the retinal vasculature manifests laterality are still unclear and are beyond the scope of the present work.

A limitation of our study is the cross-sectional design. Longitudinal studies examining retinal changes and progression of PVS and other SVD markers should assess retinal arteriolar narrowing and vessel sparseness as predictors of PVS enlargement, SVD lesion formation and cognitive decline. We were only able to obtain valid quantitative PVS and retinal measurements in a subset of the sample resulting in many subjects being excluded). However, other image analysis work in this area report similar ratio of brain imaging data that can be processed [49]. Another limitation is the unknown effect of inaccuracies in the semi-automatic measurements of the retinal vasculature (for instance, CRAE and CRVE are subject to magnification effects and refractive error; FD is dependent on the vessel segmentation accuracy [41], which in turn depends on image quality, presence of cataracts and floaters) [50].

Strengths of our study include the use of multiple computational measurements of PVS burden, the careful blinding of retinal and brain image analysis, and the inclusion of relevant risk factors and vascular disease.

## 5. CONCLUSIONS

Our study shows that older persons with narrower retinal arterioles and reduced branching complexity are more likely to have more and larger visible PVS. This suggests that PVS widening and increasing numbers are indeed markers of adverse microvascular health. Further studies are required to understand these mechanisms and their relation to brain fluid and waste clearance, and risk of dementia and stroke [2].

## ACKNOWLEDGEMENTS

### Funding Sources

The LBC1936 Study was funded by Age UK and the UK Medical Research Council (http://www.disconnectedmind.ed.ac.uk/) (including the Sidney De Haan Award for Vascular Dementia). Funds were also received from The University of Edinburgh Centre for Cognitive Ageing and Cognitive Epidemiology, part of the cross council Lifelong Health and Wellbeing Initiative (MR/K026992/1), and the Biotechnology and Biological Sciences Research Council (BBSRC). The work was also funded by the EPSRC grant [LB EP/M005976/1], the Fondation Leducq Network for the Study of Perivascular Spaces in Small Vessel Disease [LB 16 CVD 05], the Row Fogo Charitable Trust [MVH Grant No. BROD.FID3668413], the European Union Horizon 2020 [PHC-03-15, project No 666881, ‘SVDs@Target’], the UK Dementia Research Institute at the University of Edinburgh and the British Heart Foundation Centre for Research Excellence, Edinburgh.

## Author’s Contributions

Lucia Ballerini: development of the PVS segmentation tool and application to the cohort scans, checking the scans and editing, VAMPIRE software development, drafting and final approval the manuscript.

Maria del C. Valdés Hernández and Susana Muñoz Maniega: image analysis and final approval of the manuscript.

Sarah McGrory: VAMPIRE data collection, revision and final approval of the manuscript.

Enrico Pellegrini, Tom MacGillivray, Emanuele Trucco: VAMPIRE software development, revision and final approval of the manuscript.

Ross Henderson, Adele Taylor: data collection, revision and final approval of the manuscript. Ruggiero Lovreglio: software development, revision and final approval of the manuscript.

Mark E. Bastin: MRI protocol design and quality assurance, revision and final approval of the manuscript

Ian J. Deary: LBC chief investigator, funding, cohort recruitment, assessment, data analysis, revision and final approval of the manuscript.

Joanna M Wardlaw: study conception and design, funding, data analysis and interpretation, drafting, revision and final approval of the manuscript.

In addition, we thank the LBC1936 participants, nurses at the Edinburgh Clinical Research Facility, radiographers and other staff at the Edinburgh Imaging (https://www.ed.ac.uk/clinical-sciences/edinburgh-imaging): a SINAPSE collaboration Centre.

## Sponsor’s Role

The sponsors did not participate in the design, methods, subject recruitment, data collections, analysis or preparation of this manuscript

## Conflict of Interest Disclosure

none

## Notes

**Conflicts of interest:** none

### Competing Interest Statement

The authors have declared no competing interest.

